# WNT-driven chromosomal instability as a biomarker for PORCN inhibition

**DOI:** 10.64898/2026.04.13.718275

**Authors:** Diego Garcia-Lopez, Azedine Zoufir, Barbara Hernando, Patricia G. Santamaria, Alice Cadiz, Hector De Galard Terraube, Laura Madrid, Amy Cullen, Simon Woodcock, Ania Piskorz, Nicola Wallis, Jackie Walling, James D. Brenton, Florian Markowetz, Jason Yip, José Teles, Geoff Macintyre

## Abstract

Targeting Porcupine (PORCN), a key regulator of the WNT-signalling pathway, has shown therapeutic potential in multiple cancers. Despite strong target engagement and acceptable safety profiles through human phase I clinical trials, low phase II efficacy has stalled further clinical development. Given that aberrant WNT signalling can drive tumorigenesis by inducing chromosomal instability (CIN), we hypothesised that genomic CIN signatures might serve as a predictive biomarker to help improve response rates. Using a controlled *in vitro* model and single-cell whole-genome sequencing, we demonstrate that acute WNT-activation directly induces three distinct types of CIN: whole genome duplication, replication stress, and impaired homologous recombination. We translated these observations into a composite CIN signature biomarker that significantly correlated with both genetic dependency and pharmacological inhibition of PORCN across 195 and 24 cell lines, respectively. Through a large-scale meta-analysis of patient-derived and cell line xenografts, we established that this composite CIN signature biomarker quantitatively predicts *in vivo* PORCN inhibitor sensitivity (R=-0.71, p<0.002). By applying an optimised biomarker threshold, refined through modelling of human patient data, to the The Cancer Genome Atlas dataset, we successfully retrospectively modelled previous trial results and identified gastroesophageal cancers as a high-prevalence (36.6%) indication for future development. We validated this strategy in a mouse clinical trial of gastric and esophageal xenografts, where biomarker-guided stratification achieved an objective response rate of 60% and significantly decreased risk of progression (HR=0.21, p=0.0345). These data establish an actionable, trail-ready framework for further PORCN inhibitor clinical development.

## Introduction

The Wingless-related integration site (WNT) signaling pathway is a highly conserved group of signal transduction cascades that play crucial roles in embryonic development, tissue homeostasis, and cancer progression. This pathway regulates various cellular processes, including cell proliferation, polarity, differentiation, migration, and stem cell maintenance^1^. Dysregulation of WNT signaling has been implicated in numerous cancers^2–7^. Aberrant activation of the canonical WNT/β-catenin pathway leads to the accumulation of β-catenin in the cytoplasm and its subsequent translocation to the nucleus, where it induces the transcription of genes involved in cell proliferation and survival^1,8^. Recent research has also highlighted the role of WNT signaling in modulating the tumour immune microenvironment, influencing processes such as dendritic cell maturation, T cell differentiation, and immune cell infiltration^9^.

Given its central role in driving tumorigenesis and its well-established involvement in resistance to treatments, WNT signalling has been an appealing target for drug discovery and development in cancer^10,11^. A number of strategies and molecular targets have been explored to modulate WNT signalling for therapeutic purposes^12,13^. Some of the most clinically advanced are small molecule inhibitors of PORCN^12^. PORCN, or Porcupine, activates WNT signalling via palmitoylation of WNT ligands. Inhibiting PORCN prevents palmitoylation causing WNT ligands to be degraded in the endoplasmic reticulum (ER)^14^. Many of the clinical trials for PORCN inhibitors relied on *RNF43* loss-of-function mutations and R-spondins (*RSPO*) fusions as biomarkers to select patients for treatment. RNF43 inhibits WNT signaling by reducing the membrane level of Frizzled, WNT receptor, serving as a negative feedback mechanism^15^. Extracellular RSPO activates WNT signaling by preventing RNF43 binding to Frizzled and LRP5/6 co-receptors^16^. These trials sought to exploit the hyperactive WNT signalling driven by *RNF43* loss-of-function and *RSPO* fusions. However, despite this strong biological rationale, trials enriched for these alterations yielded only a small number of responders^17^. Consequently, the clinical development of PORCN inhibitors pivoted to immunotherapy combinations, but this also yielded only modest efficacy. Because monotherapy efficacy was observed in a subset of patients^17^, this suggests that the clinical limitation lies not with the drug, but with the biomarker. Accurately stratifying patients with a superior biomarker could rescue PORCN inhibitor monotherapy, highlighting a clinical unmet need for a more robust predictor of WNT–pathway dependency.

We have recently developed a new class of biomarkers based on signatures of chromosomal instability (CIN) that can assist with therapy selection^18,19^. This framework decodes patterns of DNA copy number change, quantifying different types of CIN based on the specific pattern that is left in the genome^18^ and has been demonstrated to be a robust predictor of resistance to chemotherapy^19^. Given emerging evidence that hyperactive WNT signalling can activate CIN^13^, we hypothesised that measuring the downstream accumulation of WNT-induced damage could serve as a superior biomarker for selecting patients for treatment with PORCN inhibitors.

## Results

### WNT-activation induces three distinct types of CIN

To establish precisely which types of CIN might be driven by WNT signalling, we developed a controlled *in vitro* system (**Figure 1a**) to induce WNT-driven CIN in a model that is CIN-tolerant, with a near normal genome background: hTERT-immortalised human retinal pigment epithelial (RPE-1), *TP53* knockout (*TP53*-/-) cell line. Cells were treated with either recombinant WNT3a ligand to activate WNT signalling or PBS as a control. Following successful confirmation of WNT pathway activation via quantitative PCR for *AXIN2* (**Supplementary Figure 1**), a direct transcriptional target of canonical WNT signalling^20^, the cells were cultured for five days to allow for the accumulation of DNA damage. We then performed single-cell whole-genome sequencing (scWGS) to enable ultra-sensitive detection of copy number alterations (CNAs) that emerge immediately following acute WNT activation.

**Figure 1.**
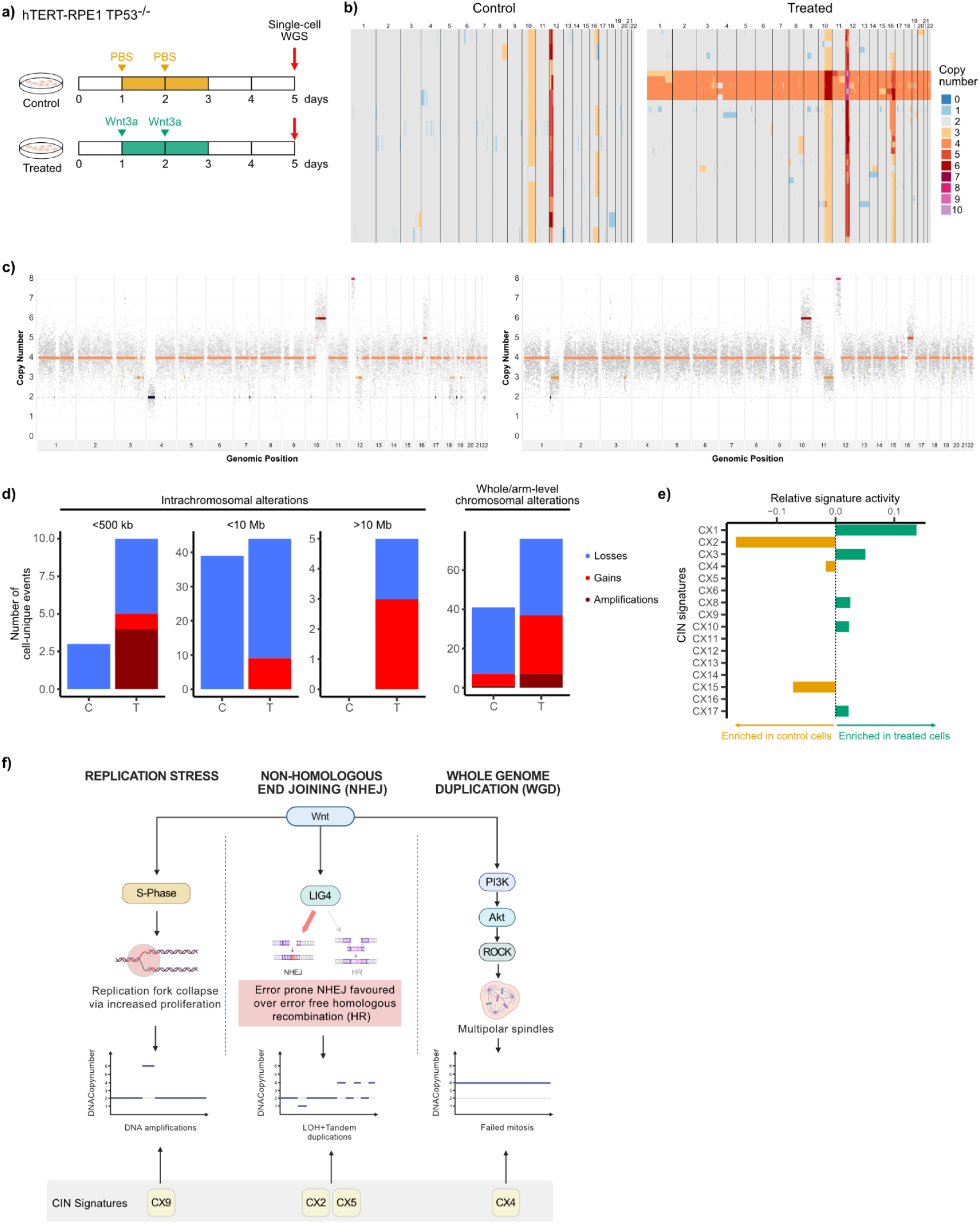
Single cell whole-genome sequencing (scWGS) of CIN tolerant, WNT-activated cell lines shows induced chromosomal instability. **a**) A schematic of the experimental pipeline for the interrogation of WNT-induced chromosomal instability. **b**) Heatmap showing single-cell copy number alterations of the control line (left) and WNT3a-treated line (right). Each row represents a single cell and each column represents a chromosome. Grey colouring represents diploid segments, blue colours represent losses, and red/purple colours represent gains. **c)** Representative absolute copy number profiles of two WNT-induced cells included in b. Coloured horizontal lines indicate copy number segments, while grey dots indicate bins. **d)** Barplots showing the number of cell-unique copy number alterations in the control (C) and the treated (T) cells based on size and chromosomal location. Alterations are classified as losses, gains or amplifications, and bars coloured accordingly. **e)** Barplots showing relative enrichment of signature activities in control (yellow) and treated (green) cells. **f)** A schematic showing how WNT-signalling can drive chromosomal instability and how CIN signatures can be used as a readout of these CIN types. Created with BioRender.com.

Analysis of the scWGS data revealed distinct patterns of copy number aberrations (CNAs) unique to the WNT3a-treated population. We observed a distinct subset of cells that had undergone whole genome duplication (WGD) (**Figure 1b**), an event entirely absent in the PBS-treated controls. Compared to the CNAs that accumulated in the control cell population, which were typical for *TP53-/-* cells in culture^21^, the WNT-activated cells showed an increase in short (<500kb) amplifications (**Figure 1d**), which are highly indicative of replication stress-induced DNA damage^22^, and an increase in intrachromosomal gains that point toward impaired homologous recombination^23,24^. Some individual cells showed evidence of all three types of genomic events (for example **Figure 1c**).

We next sought to translate these induced CNAs into our standardised CIN signature framework^18^. Using our recently developed single-event CIN signature mapping approach^21^, we evaluated the specific genomic patterns of the induced CNAs, limiting our analysis to cell-unique events to guarantee quantification of ongoing, *de novo* CIN rather than historical clonal alterations. This analysis revealed a clear induction of impaired homologous recombination (IHR) biology, captured predominantly by signature CX3, alongside replication stress (RS), captured by signature CX8 (**Figure 1e**). It is important to note that, because these acute events were induced over a short 5-day window in an otherwise stable, near-diploid genomic background, the full, complex pan-cancer signatures representing these biologies (such as WGD:CX4, IHR:CX2, CX3, CX5, and RS:CX8, CX9 and CX13) may require chronic WNT activation and long-term clonal selection to fully manifest. Nevertheless, the foundational biological processes driving these specific CIN phenotypes appear to be demonstrably active immediately following WNT stimulation.

Based on these empirical observations and existing literature, we propose a working model wherein aberrant WNT signalling drives CIN predominantly through three distinct mechanisms (**Figure 1f**). First, WNT signalling can drive early S-phase entry and exit, leading to replication fork collapse, with subsequent DNA amplification, characteristic of replication stress^25–27^. Second, because β-catenin directly targets DNA ligase IV (LIG4), a key enzyme in the non-homologous end joining (NHEJ) pathway, activated WNT signalling favours double strand break repair via error-prone NHEJ^28^. This in effect creates a BRCA-ness phenotype and accumulation of DNA losses and gains reminiscent of homologous-recombination deficiency^29^. Finally, WNT signalling, via cross talk with elements of the PI3K/AKT pathway^3,8,30^, can also promote tolerance to WGD^31^.

### PORCN dependency correlates with CIN signature levels *in vitro*

Having established that WNT activation can induce specific types of CIN - namely WGD, IHR and replication stress - we sought to identify an optimal combination of CIN signatures^18^ representing these biologies to serve as a predictive biomarker for PORCN inhibition. To achieve this, we leveraged genetic perturbation data from the Cancer Dependency Map (DepMap)^32,33^. We systematically evaluated all combinations of signatures linked to IHR (CX2, CX3, CX5), WGD (CX4) and replication stress (CX8, CX9, CX13), for correlation with PORCN knockdown (KD) dependency (Demeter2 v6) across 195 cell lines derived from 29 distinct tumour types. For each evaluated combination, a composite score was computed as the sum of z-score normalised, non-negative signature activities (see **Methods**). To identify the most robust predictor, we ranked these combinations using a Mann-Whitney U test comparing groups split at the median composite score (**Supplementary Table 1**). The top-performing composite score, consisting of CX2, CX4, CX5, and CX9, demonstrated a significant correlation with response to PORCN KD (Kendall’s tau=-0.143, p-value=0.001; **Figure 2a**), indicating that higher signature activities associate with a stronger impact on cancer cell line viability.

**Figure 2.**
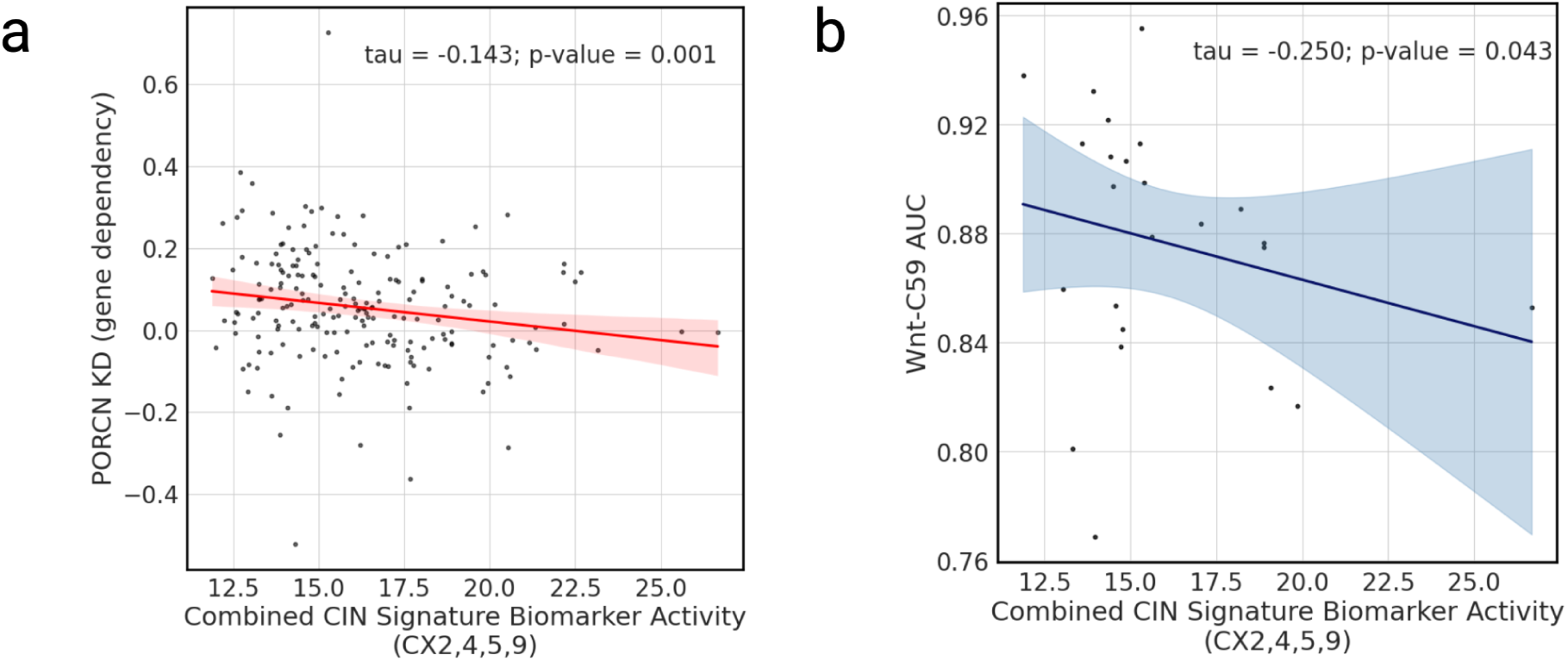
Composite CIN signature biomarker and PORCN knockdown/inhibition. **a)** Scatter plot showing correlation between combined CX2, CX4, CX5 and CX9 CIN signature activity and dependency to knockdown of PORCN across 195 cell lines. The lower the gene dependency score, the stronger the effect of knocking down the gene. **b)** Scatter plot showing correlation between combined CX2, CX4, CX5 and CX9 CIN signature activity and response to inhibition of PORCN with WNT-C59 across 24 cell lines. AUC represents the area under the dose response curve, with lower AUC indicating sensitivity. Each dot represents a cell line, and correlations were assessed using Kendall’s rank correlation coefficient (tau). P-values are from a one-tailed test.

To orthogonally validate the quantitative relationship between the composite CIN signature biomarker and PORCN dependency, we evaluated pharmacological inhibition data for the small molecule PORCN inhibitor WNT-C59, also sourced from DepMap. Because DepMap screens rely on standardised *in vitro* culture conditions that lack exogenous WNT ligands, the vast majority of cell lines do not rely on WNT signalling and are inherently insensitive to PORCN inhibition. Therefore, we restricted our analysis to a subset of sensitive models (IC50<10μM, GDSC2, n=24 cell lines). This filtering isolates a biologically relevant cohort of cell lines that likely harbour endogenous WNT activation, such as autocrine WNT signalling^34^. Within this enriched group, we observed a correlation between higher biomarker activities and increased sensitivity to WNT-C59 (Kendall’s tau=-0.25, p-value=0.043; **Figure 2b**). Taken together, these data indicate that WNT inhibition via PORCN is a viable therapeutic strategy for cancer cells with CIN, and that our rationally designed, multi-signature CIN biomarker can capture this WNT-driven vulnerability.

### PDX meta-analysis defines a PORCN biomarker threshold

Historically, PORCN inhibitors have progressed to clinical trials based on *in vivo* proof-of-concept studies using *RSPO/RNF43* alterations as predictive biomarkers^35–41^. Because these studies were primarily designed to demonstrate therapeutic efficacy, they were statistically underpowered to assess biomarker efficacy (typically 1-5 models per arm). Despite strong mechanistic support behind RSPO/RNF43 status being a predictor of PORCN inhibitor efficacy, collectively, these data did not validate this premise. There were multiple documented instances of both false positives (e.g. *RNF43* mutant cell line model CAPAN2 showing resistance^39^) and false negatives (e.g. wild-type VCAP showing sensitivity^41^). This potentially explains the subsequent modest efficacy seen in human trials selection of patients based on *RSPO/RNF43* alterations. However, collectively, this quantity of existing *in vivo* data does provide an opportunity to achieve robust statistical power for biomarker analysis, therefore, rather than conduct another inherently underpowered small-scale study to validate the relationship between our composite CIN signature biomarker and treatment response *in vivo*, we performed a large-scale meta-analysis of existing *in vivo* data.

To overcome the heterogeneous landscape of these historical studies, we compiled data from 28 sources, encompassing 123 distinct patient-derived or cell line xenograft (PDX or CDX) models and 5 PORCN inhibitors across 174 experimental regimens^35–43^, and applied a stringent quality control and standardisation pipeline (**Figure 3a**, see **Methods**). To ensure clinical translatability, we restricted our analysis to doses that exhibited drug exposure levels equivalent to those achieved in human clinical trials. We determined the minimum efficacious dose (MED) for each drug by aligning the minimum blood plasma concentration (*C_min_*) in mice with the steady-state *C_min_* from human patients treated at recommended doses^34,38,44–48^ (**Supplementary Figures 2-5, Supplementary Tables 4-6**). By excluding models with non-translatable pharmacokinetic profiles and harmonising efficacy endpoints to a fixed 17-day tumour inhibition (T/C) ratio, we mitigated study-to-study variation. Consistent with established benchmarks^49,50^, models with a T/C ratio <50% were classified as sensitive. Following this filtering, we retained a high-confidence cohort of 15 PDX/CDX models (predominantly gastrointestinal) from 6 studies, representing a combined total of 280 mice (**Supplementary Table 2**). By computing the composite CIN biomarker from the genomic data of these specific models, we established a highly controlled dataset to directly assess the predictive power of our CIN signature and RSPO/RNF43 biomarkers.

**Figure 3.**
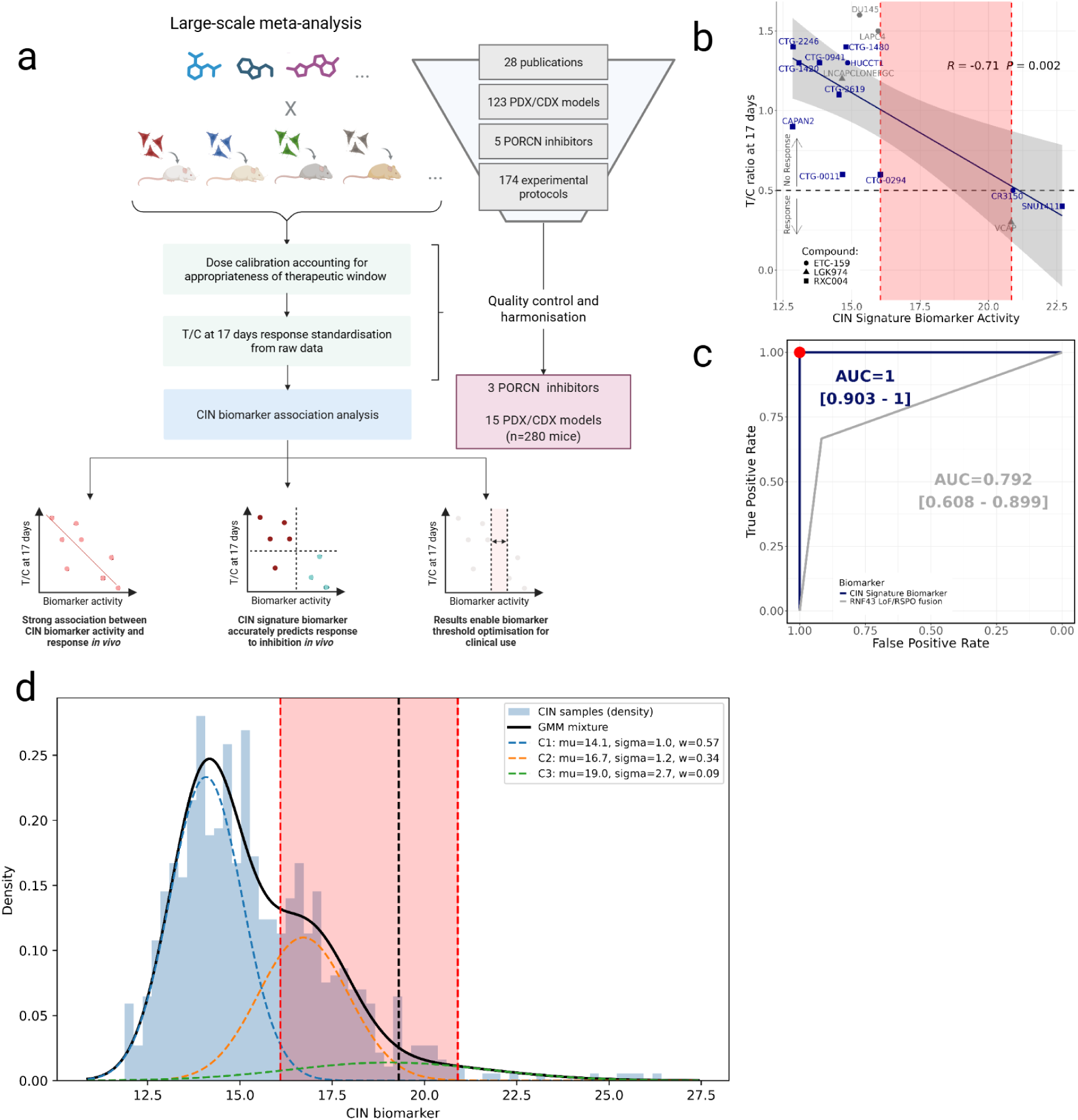
Large-scale meta-analysis of PORCN inhibitor response and association with CIN signatures in vivo. **a)** Schematic representation of large-scale meta-analysis of patient derived xenografts treated with PORCN inhibitors. **b)** T/C ratio versus composite CIN signature biomarker activity (CX2, CX4, CX5 and CX9). Each dot represents a PDX/CDX, with blue colouring indicating gastrointestinal tumours. Pearson’s correlation with one-sided test. Red dashed lines and shaded area represent the interval for an optimal predictive threshold for clinical implementation. **c)** Receiver operator characteristic curves for predicting PDX PORCN inhibitor response for the combined CIN signature biomarker and RNF43/RSPO fusion biomarker. AUC Clopper–Pearson CI95% is represented in brackets. AUC = Area under the curve. **d)** Density plot showing composite CIN signature biomarker activities for 766 cholangiocarcinomas, pancreatic, prostate and colorectal cancers in TCGA. The low, intermediate and high fit Gaussian distributions are indicated by blue, yellow and green dotted lines, respectively. The red shaded region corresponds to the optimal biomarker interval from panel b). The black dotted line is the optimal threshold which represents the Bayes Decision Boundary between the intermediate and high biomarkers groups.

The composite CIN signature biomarker demonstrated significant negative correlation with 17-day T/C ratio (Pearson R=-0.71, p-value=0.002, **Figure 3b**). This confirms that higher CIN signature activity quantitatively predicts greater sensitivity to *PORCN* inhibition. Overall predictive performance showed near-perfect classification accuracy (AUC=1.0; 95% CI: 0.903-1.00), while the standard RNF43 LoF/RSPO fusion biomarker showed markedly lower performance (AUC=0.792; CI:[0.608-0.899]) in the same cohort (**Figure 3c** and **Supplementary Figure 6**). These data provide *in vivo* evidence that decoding WNT-driven chromosomal instability offers a superior predictive framework for PORCN inhibitor patient selection compared to upstream genetic alterations.

Having established the predictive capacity of the continuous composite CIN signature biomarker, we next sought to define an optimal, binary cut-off for prospective clinical implementation. To position this biomarker for a future Phase II clinical trial, it is critical to maximise the positive predictive value (PPV) - equating to the selected objective response rate (ORR) - to ensure trial feasibility and patient benefit. While our PDX meta-analysis showed a perfect retrospective separation between sensitive and resistant models, the data was sparsely distributed at the critical decision boundary. Specifically, we observed a wide activity window between the highest-scoring resistant model (biomarker activity ∼16) and the lowest-scoring sensitive model (biomarker activity ∼21) (**Figure 3b**). Relying solely on this sparse *in vivo* data to define a precise clinical threshold risked an arbitrary boundary selection. To resolve this uncertainty and establish a biologically robust threshold, we leveraged large-scale human genomic data from the TCGA. We isolated tumour types with documented preclinical and clinical responses to PORCN inhibitors (pancreatic, biliary, prostate and colorectal cancers) and applied Gaussian mixture modelling to the composite CIN signature biomarker density distribution. To define a statistically robust threshold within our predefined activity window (16.1 to 20.9), we modelled the biomarker distribution as three distinct states representing low, intermediate and high biomarker activity (**Figure 3d**). We established the optimal clinical cutpoint at the Bayes Decision Boundary separating the intermediate and high biomarker distributions (t ≥ 19.33). By integrating *in vivo* validation boundaries with human population-level density modeling, this approach translates a quantitative biomarker score into a highly actionable, trial-ready clinical classifier.

### CIN signature prevalence identifies gastric and esophageal cancers as high-priority indications

To assess the utility of the optimised biomarker threshold, we aimed to retrospectively predict the outcomes of recent Phase I/II PORCN inhibitor trials for RXC004/Zamaporvint, ETC159 and LGK974. These trials demonstrated limited efficacy, even when enriching for standard *RNF43/RSPO* alterations. Because primary tumour material from these patients was unavailable for direct CIN signature profiling, we leveraged TCGA^51^ to estimate the baseline prevalence of biomarker positive patients (combined signature activity ≥ 19.33) across the specific indication evaluated in these historical trials. Using these population prevalences, we calculated a predicted Objective Response Rate (pORR) and the probability of achieving an ORR >30% (a standard benchmark for continued clinical development) to determine if our biomarker could explain these historic trial outcomes.

Our TCGA-derived predictions aligned well with the observed response rates in monotherapy trials^17,52^ (**Figure 4a**) and combination trials^45,53^ (**Supplementary Figure 7**, PORCN inhibitors plus anti-PD1 immunotherapy) where we applied the Bliss Independence model^54^ to mathematically infer the PORCN inhibitor-driven component of the clinical response (see **Methods**). Cohorts with low predicted biomarker prevalence, such as colorectal cancer (2.9%), showed correspondingly low clinical benefit (0% ORR). Conversely, the combined ovarian and endometrial cancer cohorts exhibited predicted biomarker prevalence of 23.9% and 21.7% respectively, yielding a combined pORR of 22.2%, which exactly matched the observed combination ORR (22.2%)^45^. Similarly, our biomarker also predicted the combination ORR (12.5%) in both the NCT02521844 ETC159 trial in solid tumours^45^, and the NCT01351103 dose expansion trial evaluating LGK974 in cutaneous melanoma^55^ (**Supplementary Figure 7)**. By successfully predicting the outcomes of these historic trials, we confirmed that our quantitative threshold translates robustly to human populations. Furthermore, these data suggest that the modest efficacy observed in previous trials may have been influenced by indication selection based on RNF43/RSPO biomarkers, providing a clear rationale to use our CIN signature to systematically identify a high-prevalence, optimal indication for future clinical development. An analysis of the TCGA pan-cancer cohort (n=7,660 MSS tumours) revealed that *RSPO3* fusions and *RNF43* loss-of-function mutations were rare events (10 and 22 samples, respectively) and are almost entirely absent among CIN biomarker-predicted responders, with only 3 of 979 responders (0.3%) harbouring either alteration (**Supplementary Table 9**). In colorectal cancer specifically, where these alterations are most prevalent, none of the seven *RSPO3* fusion-positive or three *RNF43* loss-of-function patients were predicted responders, demonstrating that the CIN biomarker identifies a distinct, non-overlapping patient population not captured by existing WNT pathway genomic markers.

**Figure 4.**
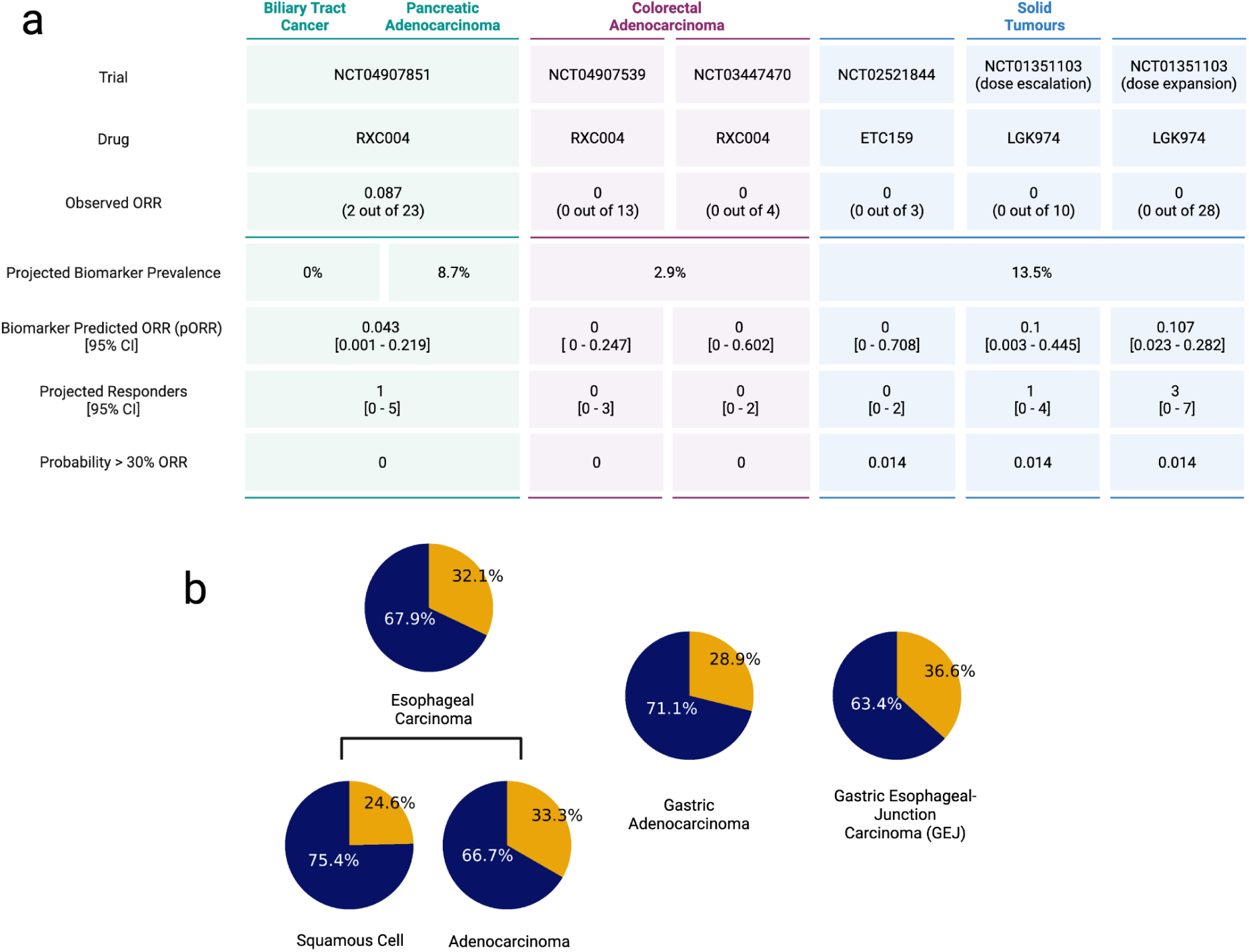
Monotherapy trial outcome prediction and optimal indication identification using CIN signature biomarker. **a)** CIN signatures predicted a responder prevalence of 0% for Biliary Tract Cancer, 8.7% for Pancreatic Adenocarcinoma, 2.9% for Colorectal Adenocarcinoma, and 13.5% for Solid Tumours. In the NCT04907851 Biliary Tract Cancer cohort (RXC004, n=23), the observed Objective Response Rate (ORR) was 0.087 (2/23), corresponding to the predicted ORR of 0.13 (3/23, 95% CI: [0.028–0.336]). The observed ORR was 0 across all remaining clinical trials, including Pancreatic Adenocarcinoma (NCT04907539, n=13), Colorectal Adenocarcinoma (NCT03447470, n=4; NCT02521844, n=3), and Solid Tumours (NCT01351103 LGK974 dose escalation, n=10; dose expansion, n=28). For the NCT01351103 Solid Tumours trial using LGK974, the observed ORR of 0 contrasts with higher predicted ORRs of 0.1 (1/10) and 0.107 (3/28) for the escalation and expansion cohorts, respectively. **b)** Predicted response rates in esophageal, gastric and gastroesophageal junction (GEJ) cancers based on TCGA cohort analyses.

Having established the biomarker’s translational potential, we systematically evaluated TCGA to identify an optimal indication for a future biomarker-enriched Phase II trial. We defined an optimal indication as one characterised by high predicted biomarker prevalence (enabling rapid trial recruitment) and high-unmet clinical need (with the potential to satisfy fast-track and accelerated approval criteria). This analysis revealed several high-prevalence indications (**Supplementary Table 7**), including breast cancer, sarcomas and upper gastrointestinal cancers. Among these, a combined indication of esophageal, gastric and gastroesophageal junction (GEJ) tumours emerged as the ideal candidate (**Figure 4b**). These cancers are fundamentally driven by CIN^56^, frequently present at advanced stages where standard-of-care therapies offer limited survival benefit^57^ and represent a profound global unmet-need^58^.

To stress-test the feasibility of a trial in this combined gastroesophageal indication, we modeled projected outcomes using a highly conservative performance estimate (see **Methods**). Applying the lower bound of the 90% confidence interval for biomarker PPV observed in our PDX meta-analysis (36.8%), we simulated a hypothetical 30-patient trial arm. Even under this worst-case statistical scenario, the model estimates a 85.5% probability of achieving an ORR > 30%. This projection demonstrates a high likelihood of clinical success when deploying the CIN biomarker in this specific patient population.

### Prospective biomarker validation in a gastric/esophageal mouse clinical trial

To assess the clinical utility of our composite CIN signature biomarker in our identified optimal indication, we conducted a prospective mouse clinical trial (MCT) using PDX models of gastric and esophageal cancer (**Supplementary Table 10**). The study utilised a parallel-arm design, evaluating the clinical-stage PORCN inhibitor RXC004 (Zamaporvint) administered at 5 mg/kg once daily (QD) in the treatment arm (n=20 models) compared to an untreated vehicle control arm (n=27 models). Models were enrolled without prior biomarker selection (an “all-comers” approach) to allow for an unbiased post-hoc assessment of biomarker specificity. The primary clinical objective of the trial was to demonstrate that biomarker stratification could significantly enrich the PPV, equating to a higher selected ORR. The trial was powered to target an expected PPV of 50% (with a 90% confidence interval and a target precision of +/- 30%). Assuming a baseline biomarker positivity prevalence of 36%, this design necessitated a minimum of 20 treated models. Following enrollment and baseline genomic profiling, our predefined optimal threshold prospectively stratified 5 of the 20 treated models into the predicted sensitive group (**Figure 5a**). Systemic toxicity was monitored via body weight measurements daily and clinical observations weekly. In the treatment group, four animals (20%) were euthanized prior to the study completion due to reaching humane endpoints. Three were euthanized due to body weight loss exceeding 20% of baseline. Of these, one was euthanized due to disease-related cachexia; this event was considered unrelated to Zamaporvint administration as the animal displayed a high tumour burden (>1800 mm^3^). The remaining mouse was terminated due to the development of repeated ascites. One additional mouse passed away before reaching the study endpoint and was not attributed to a treatment-related toxicity. No other significant clinical signs of distress were observed in the remaining cohort, and all other animals successfully completed the study.

**Figure 5.**
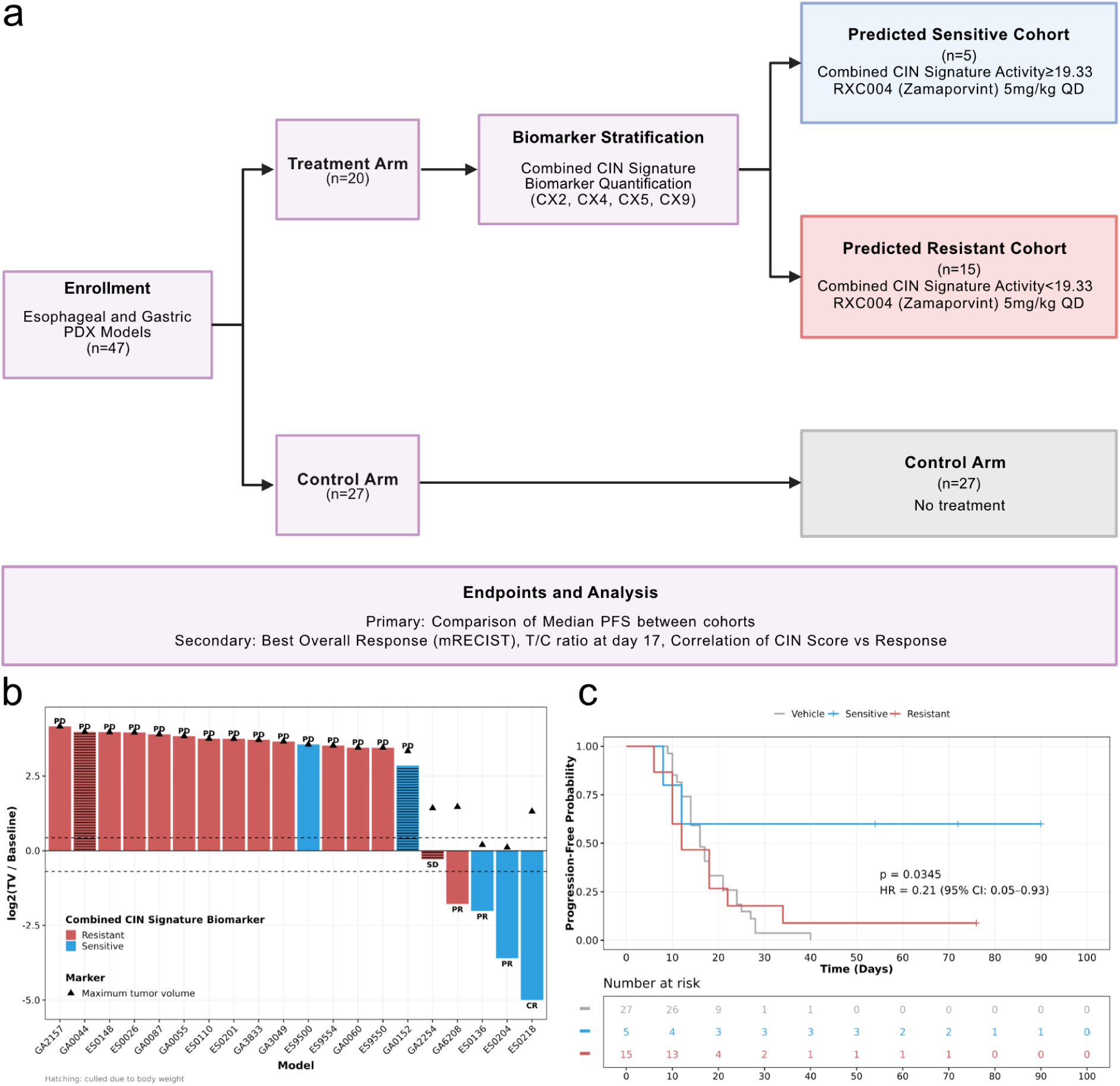
Results of a PORCN inhibitor mouse clinical trial. **a)** Schematic of the mouse clinical trial design. **b)** Waterfall plot of the tumor volume at endpoint relative to the baseline for all mice treated with RXC004. Black dotted lines show progressed disease and partial response. Peak tumor volume reached during the study is represented as a ▲. mRECIST responses at endpoint are specified for each sample: Complete Response (CR), Partial Response (PR), Stable Disease (SD) and Progressed Disease (PD). **c)** Progression Free Survival (PFS) curve comparing the different cohorts: Combined CIN Signature Biomarker sensitive (blue) Combined CIN Signature Biomarker Sensitive resistant (red) and control group (gray).

To evaluate therapeutic efficacy, we assessed tumour volume changes using modified murine RECIST (mRECIST) criteria (see **Methods**). Across the unselected treated cohort, we observed one complete response (CR) and three partial responses (PR), yielding a baseline ORR of 20% (4 out of 20). Biomarker-guided stratification successfully enriched for therapeutic benefit, more than doubling the expected efficacy to achieve a selected ORR of 60% within the predicted sensitive group (3 out of 5) (**Figure 5b**). The ORR in the predicted resistant cohort was 6.67% (1 out of 15) and 0% in the control group. Furthermore, the biomarker successfully identified 14 of the 15 non-responding models as predicted resistant, achieving a specificity of 93.33% and demonstrating its utility in preventing the administration of ineffective therapies. This diagnostic enrichment translated into a substantial survival advantage. Kaplan-Meier analysis of progression-free survival (PFS) demonstrated no significant difference between the predicted resistant models and the untreated vehicle control, highlighting the clinical futility of treating biomarker-negative tumours (Hazard Ratio (HR)=0.99, p-value=0.97). In contrast, the biomarker-selected, predicted sensitive cohort exhibited a significantly decreased risk of progression when treated with RXC004 (HR=0.21 [95%CI: 0.05-0.93], p-value=0.0345). After a median follow-up of 42.75 days, progression events occurred in 40%, 93.33% and 100% of mice in the predicted sensitive, resistant and control cohorts, respectively (**Figure 5c**). Together, these prospective *in vivo* data validate our gastroesophageal outcome modeling and demonstrate that the composite CIN signature identifies a subpopulation that derives survival benefit from PORCN inhibition.

To further characterise the depth and durability of therapeutic benefit within the biomarker-selected cohort, we longitudinally tracked the two models that achieved the best responses. In model ES0218, RXC004 treatment induced complete macroscopic tumour eradication (0 mm³), whereas model ES0204 regressed to a state of minimal residual disease (13 mm³). To evaluate the persistence of these deep responses, both models were subjected to a treatment holiday by withdrawing the PORCN inhibitor (**Supplementary Figure 10**). The complete response in ES0218 proved durable, with no evidence of tumor regrowth observed after 30 days off treatment. In contrast, model ES0204 exhibited tumour recurrence 10 days post-withdrawal. Upon rechallenging this model with the original RXC004 dosing regimen, the tumour rapidly regressed back to its prior minimal residual volume. These individual responses highlight the capacity of the CIN signature to identify tumours with exceptional sensitivity to PORCN inhibition. Furthermore, the successful re-induction of regression in ES0204 suggests that acquired resistance does not immediately emerge during brief treatment interruptions, indicating that planned drug holidays and subsequent rechallenge could serve as a viable and effective clinical management strategy for biomarker-selected patients.

## Discussion

We have previously established the utility of CIN signatures as predictive biomarkers for standard-of-care chemotherapies across multiple different tumour types^19^. Here, we extend this paradigm to targeted therapies, specifically addressing the significant clinical gap in treating WNT-driven cancers. While PORCN inhibitors have been shown to effectively suppress WNT signaling and exhibit tolerable clinical safety profiles, their clinical development has not progressed beyond Phase II trials.

A recognised limitation in evaluating new biomarkers in this space is the scarcity of primary tumour material from historic PORCN inhibitor trials, precluding direct retrospective validation. To overcome this, we employed a robust translational modeling approach. By mapping our optimised diagnostic threshold onto TCGA cohorts, we successfully explained the limited efficacy of past unselected trials and systematically identified gastroesophageal cancers as an optimal, high-prevalence clinical indication. We validated this strategy using an unselected mouse clinical trial, where biomarker stratification successfully enriched the ORR to 60% and conferred a progression-free survival advantage. Furthermore, the observation of durable complete responses and successful rechallenge following drug holidays in our MCT suggests that CIN-stratified patients could benefit from flexible dosing regimens.

Looking forward, the clinical implications of this framework extend beyond gastroesophageal cancers. Given the fundamentally tissue-agnostic nature of chromosomal instability, this composite CIN signature biomarker provides a strong rationale for pan-cancer basket trial designs, enabling the broader development of PORCN inhibitors across diverse solid tumours. Furthermore, the methodological pipeline established here, linking a specific oncogenic driver to its characteristic CIN phenotype to build a predictive diagnostic, could be systematically applied to rescue other targeted therapies that have stalled due to poor patient selection. Ultimately, the successful validation of CIN signature biomarkers in prospective clinical trials could pave the way for a new paradigm in precision oncology, matching patients to therapies based on the functional scars left within their genomes.

## Methods

### Induction of WNT-driven chromosomal instability

#### Cell line model and cell culture

The hTERT-immortalized normal human retinal pigment epithelial (RPE-1) cell line expressing Cas9 was a kind gift from Dr. Felipe Cortés-Ledesma (CNIO, Madrid, Spain). The hTERT RPE-1 Cas9 *TP53^-/-^* cell line was generated as previously described^21^ and cultured in Dulbecco’s Modified Eagle Medium Nutrient Mixture F12 (DMEM F12; Sigma Aldrich) supplemented with 10% fetal bovine serum (FBS; Sigma Aldrich), 1% penicillin/streptomycin (Pen/Strep; Solmeglass) and 1% L-glutamine (Gibco) and grown at 37°C and 5% CO_2_. WNT3a (R&D Systems) was added to cells (200 ng/ml) and refreshed after 24 h.

#### RNA isolation, reverse transcription PCR and gene expression analysis

Total RNA from control (PBS) and WNT3a-treated hTERT RPE-1 Cas9 *TP53^-/-^*cells was extracted using the RNeasy mini kit (Qiagen #74004). 1,5 µg or RNA was transcribed using QuantiTech RT kit (Qiagen # 205311). Quantitative PCR was performed using SYBR Green PCR Master Mix (Applied Biosystems #2504552) and *AXIN2* primers (forward 5’-TACCGGAGGATGCTGAAGGC-3’; reverse 5’-CCACTGGCCGATTCTTT-3’). Gene expression was normalised to *PPIA* internal control gene expression (forward 5’-ATGGTCAACCCCACCGTGT-3’; reverse-5’-TCTGCTGTCTTTGGGACCTTG-3’).

#### Single-cell whole-genome sequencing (scWGS)

Control and WNT3a-treated hTERT RPE-1 Cas9 *TP53^-/-^* cells were single cell sorted using an image based piezo-electric nanoliter dispenser (cellenONE, Scenion) in 384-well PCR plates (Eppendorf Cat. no.: 0030129504) based on cell line specific features^21^. Then, single-cell DNA whole-genome sequencing libraries were performed using a modified version^21^ of the Direct Library Preparation Plus protocol previously described (mDLP)^59^.

#### Generating single-cell copy number profiles

Reads were aligned as single-end against the human genome assembly GRCh37 using BWA-MEM (v0.7.17). Duplicate reads were then identified and marked using samtools-markdup (v1.15). To infer copy number profiles from single-cell DNA sequencing data, we defined 100 kb fixed bins along the genome. The QDNAseq R package^60^ was used to count reads per bin, which were corrected by sequence mappability and GC content using LOESS regression. DNACopy^61^ was then used to perform segmentation using default parameters. Cell bams were downsampled to 15 reads per bin per chromosome copy, and those not meeting this sequencing depth were excluded from downstream analyses. After the segmentation step, we inferred absolute copy number values across a ploidy range (1.5 < ploidy < 8) and a purity fixed at 1. The solution with the minimum RMSD was also considered as the optimal. All solutions were manually inspected to either confirm or correct for more appropriate fits. Finally, cells with excessive noise or overfragmented profiles were also excluded for downstream analyses. The filtering criteria was previously described^21^, and followed recommendations from previous studies^62,63^.

#### Exploring induced copy number alterations

We used cell-unique copy number alterations (e.g. alterations present in a cell with distinct copy number value compared to all other cells in the model) as a proxy of alterations recently generated by an ongoing CIN type. To detect cell-unique alterations, we developed a statistical approach that ensures robustness of the identified unique events by comparing the observed copy number deviations of the segment from the segments of the other cells to a null distribution^21^

To identify the possible types of CIN causing cell-unique alterations in a model, we applied a probabilistic approach to assign individual copy number events to CIN signatures^21^ by adapting our signature framework^18^, allowing the analysis of alteration patterns by individual events rather than per genomic region. This new encoding space consists of three fundamental copy number features: the segment size, the difference in copy number between adjacent segments, and the breakpoints density within a 5 Mb flanking window ^21^. Each cell-unique segment was defined in this new feature space, which was then multiplied by the signature definitions to derive a segment-by-signature vector reflecting the likelihood of an individual event being attributed to a given signature. The signature with the highest probability was then identified as the most likely CIN-related process causing the given alteration. We next quantified the fraction of cell-unique events attributed to each signature in both control and treated lines. Signature enrichment was finally assessed by subtracting, for each signature, the fraction observed in the control from that in the treated line. Positive values indicate enrichment of a given signature in the treated condition.

### *In vitro* PORCN inhibition association analysis

#### Genomic data analysis

Chromosomal instability signatures were quantified from SNP6 copy number profiles using the method described in Drews et al., 2022^18^. The copy number data was obtained from ASCAT.sc (https://github.com/VanLoo-lab/ASCAT.sc) and manually curated to derive accurate copy number profiles following the methodology described in Van Loo et al.^64^, 2010. CIN signature activity values were z-scored normalised using activities from a subset of 310 pancancer cell lines extracted from DepMap. Z-scores for each signature were truncated to a distribution between -4 and 4, then shifted to a positive Z-score distribution using the observed minimum Z-score. The sum of CX2,CX4,CX5 and CX9 z-scored activities is presented as “Combined CIN Signature Biomarker Activity”.

#### PORCN knockdown cell line data

Knockdown gene dependency data was collected from DepMap^32^ portal (depmap.org/portal/) using the DEMETER2 model^33^ which aggregates data from the Broad Institute’s Project Achilles^32^, Novartis’ Project DRIVE^65^, and a large functional breast cancer cell line screen^66^. DEMETER2, is an analytical framework for analyzing RNAi screens which incorporates cell line screen-quality parameters and hierarchical Bayesian inference. The model estimates gene dependency scores which represent the effects of a gene knockdown on the viability of a cell line.

#### PORCN inhibition drug screen data

GDSC1 and GDSC2 datasets from the Genomics of Drug Sensitivity in Cancer project (see www.cancerrxgene.org/) were used to assess PORCN inhibition *in vitro*. In these screens, the lower the AUC the higher the sensitivity to the concerned drug. Sensitive cell lines were defined as proposed by GDSC. Hence, any cell line with IC50 lower than the maximum tested drug concentration was considered to be sensitive.

#### Statistical analysis

To evaluate whether CIN signature activity was associated with sensitivity to PORCN inhibition, one-tailed Kendall’s tau correlation tests were performed. This non-parametric, rank-based method was used to assess monotonic relationships between signature scores, RNAi gene dependency values and PORCN inhibition response. A one-tailed test was selected to specifically assess whether higher signature scores were associated with increased sensitivity (i.e., stronger gene dependency or response to chemical inhibition). The null hypothesis states that there was no association or a non-negative association between signature scores and gene dependency (τ ≥ 0). The null hypothesis was rejected at a significance threshold of p-value < 0.1, indicating a statistically significant negative association consistent with increased sensitivity.

### PDX meta-analysis

#### Genomic data analysis

For the majority of the PDXs, CIN Signature quantification was performed as described above, except for LAPC4, CR3150 and SNU-1411. LAPC4 WGS data was extracted from ^67^. CR3150 WGS data was obtained from CrownBio. CTG-0011, CTG-0294, CTG-0941, CTG-1420, CTG-1480, CTG-2246, and CTG-2619 WES data was obtained from Champions Oncology. CN data for both LAPC4 and CR3150 was obtained from WGS as described in Thompson et al., 2025^19^ using a bin size of 30kb. CN data for Champions Oncology PDXs was obtained from WES as described in Thompson et al., 2025^19^ using a bin size of 50kb. SNU-1411 CN data was obtained from the Copy Number Public 24Q4 from DepMap^68^ and CIN signatures were computed from the segmented data. As described above, CIN signature activity values were z-scored normalised using activities from a subset of 310 pan cancer cell lines extracted from DepMap. Then truncated to a distribution between -4 and 4 and shifted to a positive Z-score distribution using the observed minimum Z-score. The sum of CX2, CX4, CX5 and CX9 z-scored activities transformed as described above for the *in vitro* association analysis is presented as “Combined CIN Signature Biomarker Activity” in the results section.

#### Mutation data and RSPO/RNF43 mutant definition

Somatic mutations and gene fusions data was extracted from DepMap 24Q4 release^68^. We used this data to evaluate the predicted PORCN inhibition sensitivity of the different xenografts according to the *RSPO* fusions/*RNF43* LoF biomarker – the most commonly used biomarker in PORCN inhibitor clinical trials.

*RSPO* fusions were considered to confer gain-of-function (GoF), and therefore be predictive of PORCN inhibition sensitivity according to this biomarker, according to the literature^16,69,70^.

On the other hand, *RNF43* LoF was determined if the mutation (1) is likely to truncate the protein or be pathogenic and (2) it is homozygous. Clinvar (Pathogenic and Likely Pathogenic mutations), OncoKB^71^ (Loss-of-function and Likely Loss-of-function mutations appearing in the TCGA) and literature annotation was used to evaluate the truncating effect of each mutation^72–74^. Allele frequency (AF>0.85), variant allele frequency (VAF>0.85) and genotype (GT = 1|1) columns of the OmicsSomaticMutations.csv file were used to examine the zygosity states of the variants.

Xenografts harbouring *RSPO* GoF fusions or *RNF43* LoF mutations were labelled as RSPO/RNF43 mutants. These models would be predicted to be sensitive to PORCN inhibition according to the RSPO/RNF43 biomarker.

Some publications focused on this biomarker have suggested that mutations in the WNT pathway downstream RNF43/RSPO could confer resistance to PORCN inhibition^35,37^. To account for this we looked at alterations in *APC*, *AXIN1*, *CTNNB1*, *CTNNBIP1*, *EP300* and *FBXW7*. Supposedly, *APC*, *AXIN1* and *EP300* LoF would cause resistance to PORCN inhibition. In these genes, LoF mutations were defined following the same criteria as for *RNF43*. In contrast, *CTNNB1* and *CTNNBIP1* GoF mutations would confer resistance. GoF mutations in *CTNNB1* and *CTNNBIP1* were determined based on the pathogenicity of the variant assessed using ClinVar (Pathogenic and Likely Pathogenic mutations) and OncoKb (Gain-of-function and Likely Gain-of-function mutations appearing in the TCGA). For *FBXW7*, LoF was defined following the same criteria as for *RNF43* although some *FBXW7* hotspot mutations did not need to appear in homozygosity to produce a LoF variant. As a result, the homozygosity requirement was not taken into account in those cases. Consequently, *RSPO/RNF43* mutant xenografts that also had mutations downstream these genes were considered as *RSPO/RNF43* WT for the analysis.

#### Concordance between drug exposure in mice and clinical trial patients

PDXs were treated with RXC004, ETC159, LGK974 (also known as WNT974). All of these are clinical stage inhibitors (NCT04907539, NCT02521844 and NCT01351103 respectively). To assess whether the antitumor activity of PORCN inhibition seen *in vivo* might translate to human patients, we checked concordance of PORCN inhibitor drug exposure in PDXs and patients at the recommended dose.

To achieve this, we first selected the minimum efficacious dose (MED) in the reference sensitive preclinical *in vivo* models for RXC004 (SNU1411), ETC159 (CR3150) and LGK974 (VCAP) ^75^. We defined MED, as the dose with the lowest Cmin that results in a significant antitumor activity in the reference models. For ETC159, Cmin data for the doses at which CR3150 was tested was not available. Given that all these doses were administered following the same regimen (QOD), we defined as MED the lowest dose that resulted in antitumor activity. For all other models, Cmin is defined as the minimum blood plasma concentration reached by a drug during a dosing interval. Antitumor activity was measured using T/C at 17 days.

Cmin at the MED for each drug in mice was then compared to human Cmin calculated in patients treated with the established recommended dose of the drug at steady state (C1D15). Because we did not have available Cmin data for the MED for ETC159, we compared the human Cmin data to the closest dose higher than the MED for which we have Cmin data in mice (30 mg/kg). Another caveat is that, for both LGK974 and ETC159, Cmin data was computed after a single dose administration and compared to data in patients at steady state that received the drug for 15 days. Hence, there is a risk of underestimating Cmin in mice for both ETC159 and LGK974. Cmin was standardised using the free fraction of the drug in mice and humans for RXC004, ETC159 and LGK974^38,76^ (**Supplementary Table 3**). Mouse doses were retained in cases where the recommended human dose Cmin was not significantly lower than the mouse dose Cmin (RXC004: 2mg, ETC159: 16mg and LGK974: 10mg).

#### Therapeutic window assessment analysis

Following the same principle as above, we extended the analysis to all doses tested in clinical trials lower than the established recommended dose in humans and compared it to the MED dose in mice for RXC004 and LGK974. For RXC004, we used data from monotherapy trials for the 1.5 and 1 mg doses, to which we added data at these doses in combination with immune checkpoint inhibitors (pembrolizumab and nivolumab). We also used data from patients treated on a regimen following 2 weeks on RXC004 2 mg and 2 weeks off the drug for 2 mg dose. As mentioned before, we did not have Cmin data for the MED for ETC159. As an alternative, we compared the human Cmin data to mice Cmin after a single dose of 30 mg/kg. As outlined before, this may cause underestimation of Cmin in mice for both ETC159 and LGK974. All Cmin, derived from human doses lower than the recommended dose, that were not significantly lower than mouse MED Cmin, were considered as candidate doses at which we could potentially observe antitumor response using our CIN Signature Biomarker.

#### Tumour volume data

Tumour volume data from PDXs treated with PORCN inhibitors were extracted from previous studies (**Supplementary Table 2**). We used the Plotdigitizer online tool^77^ to estimate the tumour volume or drug concentration values at the different time points for each model.

We collected a total of 123 different PDXs models that were treated at different drug concentrations of 5 different PORCN inhibitors. As a result, 174 different experimental models were extracted from 28 scientific publications (papers, posters, patents and confidential information). From this initial cohort, we only selected PDXs with tumor volume measurements that were implanted subcutaneously in mice and treated for at least 17 days. Two exceptions were made to this criteria: HUCCT1 and DU145. HUCCT1 was only treated for 15 days and DU145 for 16 days. However, in both scenarios the treatment group overgrows the control, resulting in T/C ratios>1. Hence we assumed that the remaining days would not significantly change our observations.

Treatment of these PDXs was performed either by oral gavage (for most of the PDXs) or by subcutaneous injection (VCAP and LNCAPCLONEFGC). Xenografts that lacked drug exposure data in plasma were also excluded as we could not assess the feasibility of obtaining the same response in humans. Xenografts whose drug exposure was much higher compared to the recommended dose in humans, were excluded for the same reason. CDXs without copy number data had to be excluded as we could not calculate the CIN Signature Biomarker activity. In cases in which we had the same model across different drugs or drug concentrations, we prioritised those treated with RXC004 or ETC159 and those treated with the lowest dose leaving a final cohort of 15 PDXS (**Supplementary Table 2, Supplementary Figure 5**)

#### Statistical analysis

##### Antitumor activity assessment

In order to assess the antitumor activity for each xenograft, we calculated the T/C ratio at 17 days as described^78^:

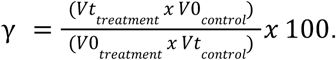

Where V_t_ is the tumor volume at time t, in this case t=17 days, and V_0_ is the tumor volume at baseline.

We fixed study end-points at 17 days to harmonise the data and control for studies heterogeneity. T/C ratios lower than 50% were considered to be indicative of sensitivity to PORCN inhibition and equivalent to a likely clinical response^49,50^.

##### Correlation between CIN Signature Biomarker and T/C at 17 days

One-tailed Pearson correlation was used to test the statistical significance of the association between T/C at 17 days and our CIN Signature Biomarker. Statistical significance was set at p-value<0.05.

##### Receiver Operator Curve comparison between CIN Signature Biomarkers and RSPO fusions/RNF43 LoF

R library pROC was used to compute the ROC curves and the AUC for both the CIN Signature Biomarker and the *RSPO* fusions/*RNF43* LoF biomarkers. The ROC curve was used to set the threshold on our CIN Signature Biomarker to predict sensitivity to PORCN inhibition.

##### Cmin comparisons between mice and humans

One-tailed Wilcoxon test was used to test whether Cmin derived from human doses were significantly lower than those Cmin observed in mice MED for RXC004. For ETC159 and LGK974, one-tailed Welch test was used for the same purpose as we did not have access to the individual raw data.

##### TCGA PORCN CIN Signature Threshold Optimization

CIN TCGA cancer samples from tumour types showing preclinical or clinical responses –pancreatic adenocarcinoma (PAAD), colorectal adenocarcinoma (COADREAD), prostate adenocarcinoma (PRAD), and cholangiocarcinoma (CHOL)-- were used to establish an optimised PORCN CIN Signature Biomarker threshold.

To identify distinct subpopulations within the CIN biomarker distribution, a Gaussian Mixture Model (GMM) was fitted to the univariate CIN biomarker values of the selected cohort using the GaussianMixture implementation in scikit-learn. The optimal number of components was selected by minimising the Bayesian Information Criterion (BIC) over a candidate range of 1-8 components. The final model was re-fitted with the optimal component count, and components were ordered by ascending mean for interpretability.

Finally, subpopulation boundaries were defined as the intersection points between adjacent weighted Gaussian component densities. The resulting thresholds represent the Bayes-optimal decision boundaries under the fitted mixture model.

### Clinical trial outcome predictions

#### Predicted Objective Response Rate (pORR)

Genomic CIN signatures were derived from 7,660 SNP array samples from the TCGA database, as per the methodology previously published^18^. The signatures were then Z-score normalised using the mean and standard deviation from the DepMap-derived CIN signature distributions (310 cell lines). Z-scores were then truncated to [-4, 4] and adding 4 to ensure a positive distribution. Samples within each cancer type in TCGA were categorised as responders or non-responders based on a threshold derived from the PDX analysis (t > 5.7). Based on the cancer type responder rates (**Supplementary Table 6**), the predicted objective response rate (pORR) was calculated as follows:

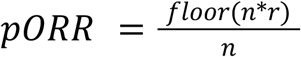

Where n is the number of patients in the trial arm, and r is the responder rate in TCGA for the specific indication in the trial. Note that only patients tested at the recommended dose (2mg for RXC004 and 16mg for ETC159) were considered for these calculations and the observed ORR calculations.

For the hypothetical gasto-esophageal trial, r is replaced by the 95% lower bound of the confidence interval on the specificity estimate (c.f. PDX analysis). This was done to estimate a conservative response rate and subsequent projected ORR. A 95% Clopper-Pearson confidence interval was also calculated when the observed ORR was greater than 0.

#### Probability of success (ORR > 30% or ORR > 50%)

Assuming the patients are selected randomly and independently from each other (which is not unreasonable given patients are distributed in different regions and trial sites), we can estimate the probability of having k responders in a trial using a Binomial distribution B(n,p) where n represents the number of patients in the trial arm/group and p represents the biomarker positive rate for the trialled indication

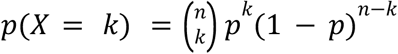

To calculate the probability of achieving more than k responders, the cumulative probability is used.

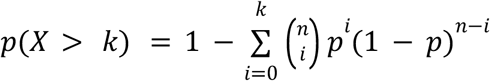

To calculate the probability of obtaining more than 30% response rate in our projected esophageal, stomach and gastroesophageal trial arm of n=20, 30 or 40 patients, we assume the response rate to be equal to the lower bound of the 95% confidence interval around the specificity estimate obtained from the PDX meta-analysis (specificity 95% CI = [0.5407-1]), therefore p=0.5407

#### Estimate monotherapy efficacy in combination therapy trials

To estimate the ORR in the case of combination therapies, the effect of the PORCN inhibitor must be inferred from the observed ORR of the combination. Assuming independence under the BLISS model of synergy^54^ (drugs in the combination act independently, at different sites of action), the ORR of the monotherapy PORCN inhibitor (ORR(PORCNi)) may be calculated from the combined efficacy with the following equation:

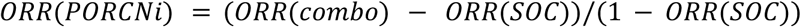

where SOC is a standard of care immunotherapy agent (e.g. Pembrolizumab or Nivolumab) given in combination to a PORCN inhibitor in the trial. In this equation, ORR(SOC) represents the lower bound 95% Clopper-Pearson confidence interval around the point estimate ORR(SOC) (**Supplementary Table 8**). This is because the point estimate for ORR(SOC) is generally larger than the observed combination ORR, as the SOC estimate is a result of being aggregated from several, highly powered trial arms. The point estimate ORR(SOC) is itself calculated as a weighted average of available monotherapy efficacy data from clinical trials for the immunological agent (weighted by the number of patients in the trial/monotherapy arm of the trial (**Supplementary Table 8**). ORR(PORCNi) was capped at 0 in case it was estimated as a negative number (due to the equation of Bliss independence above).

The predicted ORR is based on the CIN signature and therefore independent of the combined SOC/immunotherapy drug. It is estimated in the same way as in the monotherapy trials.

### Mouse Clinical Trial

#### Ethical statement and study design

All animal experiments were performed in accordance with the Institutional Animal Care and Use Committee (IACUC) of Crown Bioscience. Study protocols complied with all relevant ethical regulations regarding animal research and followed the ARRIVE guidelines. The care and use of animals were conducted in accordance with the regulations of the Association for Assessment and Accreditation of Laboratory Animal Care (AAALAC). A prospective, randomized, parallel-arm mouse clinical trial (MCT) was employed using a 1 × 1 × 1 experimental design (one animal per tumor model per treatment) to evaluate the efficacy of the PORCN inhibitor RXC004 (Zamaporvint) in gastric and esophageal cancer.

#### Animal models and husbandry

Female Balb/c nude and NOD-SCID mice, aged 6–9 weeks and weighing 19–26 g, were used for the trial. Mice were housed in groups of up to five per cage in a pathogen-free facility with a 12 h light/dark cycle, at a temperature of 20°C–26°C and 40%–70% humidity. Animals had ad libitum access to wet standard irradiated rodent chow and autoclaved filtered reverse-osmosis softened water.

#### PDX models and trial enrollment

A total of 47 patient-derived xenograft (PDX) models of gastric and esophageal cancer were utilized, including 20 models in the treatment arm and 27 models in the untreated vehicle control arm (**Supplementary Table 10**). A prospective 1 × 1 × 1 experimental design was used for treated mice in which fresh tumor fragments (approximately 2–3 mm in diameter) were harvested from stock mice and inoculated subcutaneously into the right front flank of experimental mice. Enrollment into the study occurred when the mean tumour volume reached 100–200 mm³, which was designated as Day 0. Unmatched control mice mean tumour volume data was obtained from Crown Biosciences published experiments.

#### Therapeutic intervention

Mice in the treatment group received Zamaporvint (RXC004) at a dose of 5 mg/kg via oral gavage once daily (QD). The drug was prepared weekly in a vehicle of 0.5% hydroxypropyl methylcellulose (HPMC) (w/v) in ultrapure distilled water. The dosing solution was protected from light and maintained as a cloudy, fine suspension at room temperature with continual mixing. Dosing volumes were adjusted daily based on body weight (10 µL/g). Control animals received vehicles alone as specified by the parallel-arm design.

#### Outcome measures, monitoring and termination

The primary endpoint was the change in tumor volume relative to baseline. Tumor dimensions were measured every 2 days using digital calipers, and volume (V) was calculated using the formula 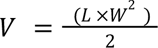, where L is the longest dimension and W is the dimension perpendicular to L.

Therapeutic efficacy was categorized using modified RECIST (mRECIST) criteria, defined by the percentage change in tumor volume (ΔV) at the study endpoint relative to the baseline volume recorded at enrollment (Day 0). Complete Response (CR) was defined as a tumor volume regression of 95% or greater (ΔV<-95%), while Partial Response (PR) was characterized by a decrease in volume between 50% and 95% (-95%<ΔV<-50%). Models demonstrating a volume change ranging from a 50% decrease to a 35% increase (-50%<ΔV<35%) were classified as Stable Disease (SD). Progressive Disease (PD) was defined as an increase in tumour volume exceeding 35% (ΔV>35%).

Body weight was monitored daily as a surrogate for systemic toxicity. Each model was terminated 24 hours post-final dose. Animals reaching humane endpoints, including body weight loss >20% relative to Day 0, tumor volume >2000 mm³, or tumor ulceration >5 mm, were euthanized via CO2 inhalation followed by cervical dislocation.

#### Statistical analysis

Survival curves were estimated using the Kaplan-Meier method and compared between biomarker-stratified groups using a log-rank test. Risk of progression was assessed using Hazard Ratios (HR) derived from univariate Cox proportional hazards models. Biomarker predictive power was evaluated by constructing contingency tables to calculate sensitivity, specificity, and PPV, with statistical significance determined by Fisher’s exact test.

## Acknowledgements

We would like to acknowledge the guidance and support of Ruth Plummer and Steven P. Jackson. We acknowledge the support of Tailor Bio. B.H., A.Ca., P.G.S. and G.M. are hosted by the Centro Nacional de Investigaciones Oncológicas (CNIO), which is supported by the Instituto de Salud Carlos III and recognized as a ‘Severo Ochoa’ Centre of Excellence (ref. CEX2019-000891-S) by the Spanish Ministry of Science and Innovation (MCIN/AEI/10.13039/501100011033). B.H., A.Ca., P.G.S. and G.M. were supported by Spanish Ministry of Science and Innovation grants PID2019-111356RA-I00 and PID2022-137042OB-I00 (MCIN/AEI/10.13039/501100011033) and co-funded by the European Regional Development Fund (ERDF-EU). B.H. was supported by philanthropists via the ‘Amigos/as del CNIO’ Programme, and also by La Caixa Foundation (ID 100010434; LCF/BQ/PR23/11980033). We acknowledge funding and support from Cancer Research UK, and the Cancer Research UK Cambridge Centre (grant nos. 22905 and 100005 to A.M.P. and J.D.B.) and the CRUK Innovation Prize PO 1121956 to G.M., A.M.P. and J.D.B. Parts of this work were funded by CRUK core grant C14303/A17197, A19274 (F.M. lab).

## Author contributions

D.G.-L. and A.Z. contributed equally to this work. G.M. conceived and designed the study. D.G.-L., A.Z., B.H., H.D.G.T, L.M., and J.T. developed the methodology and software of the study. A.Ca., A.C. and P.G.S. designed and performed in vitro experiments. D.G.-L., A.Z., B.H., P.G.S, A.Ca., H.D.G.T, L.M., A.C., S.W., N.W., J.W., J.T., F.M., J.Y., J.D.B., A.M.P. and G.M. provided access to data and/or contributed to gathering, processing and curating data. D.G.-L., A.Z., B.H., P.G.S, H.D.G.T, L.M., J.Y. and G.M. wrote the paper. G.M supervised the project. All authors had access to all of the data in the study. All authors contributed to the review and the editing of the paper. All authors approved the paper before submission.

