## Supplementary material for "WNT-driven chromosomal instability as a biomarker for PORCN inhibition"

### Supplementary Figures

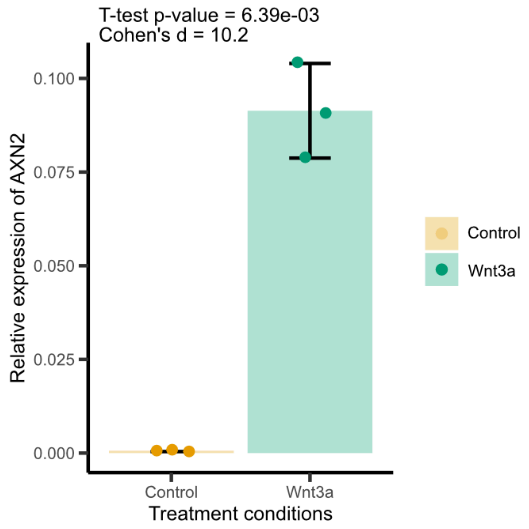


***Supplementary Figure 1: AXIN2*** **expression upon Wnt3a treatment.** *Barplot showing the relative expression of the AXIN2 mRNA in the hTERT-RPE1 TP53-/- cell line treated with PBS (Control, yellow) or WNT3a (green) determined by qPCR. The bar represents the mean value of three replicates (dots), and the error bars indicate the mean ± standard deviation.*

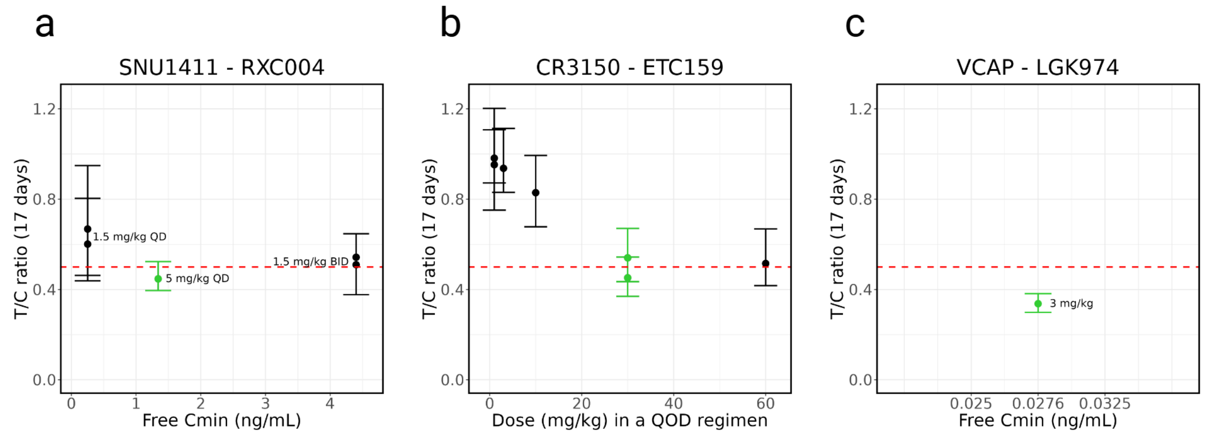


***Supplementary Figure 2: MED selection in reference models.*** *T/C ratio at 17 days compared to free Cmin (RXC004 and LGK974) or dose (mg/kg) in QOD regimen (ETC159). The red dotted line represents the threshold for antitumor activity based on T/C. Doses that result in a T/C lower than 0.5 are considered efficacious. The efficacious dose with the lowest Cmin for each drug-model combination was defined as the MED.*

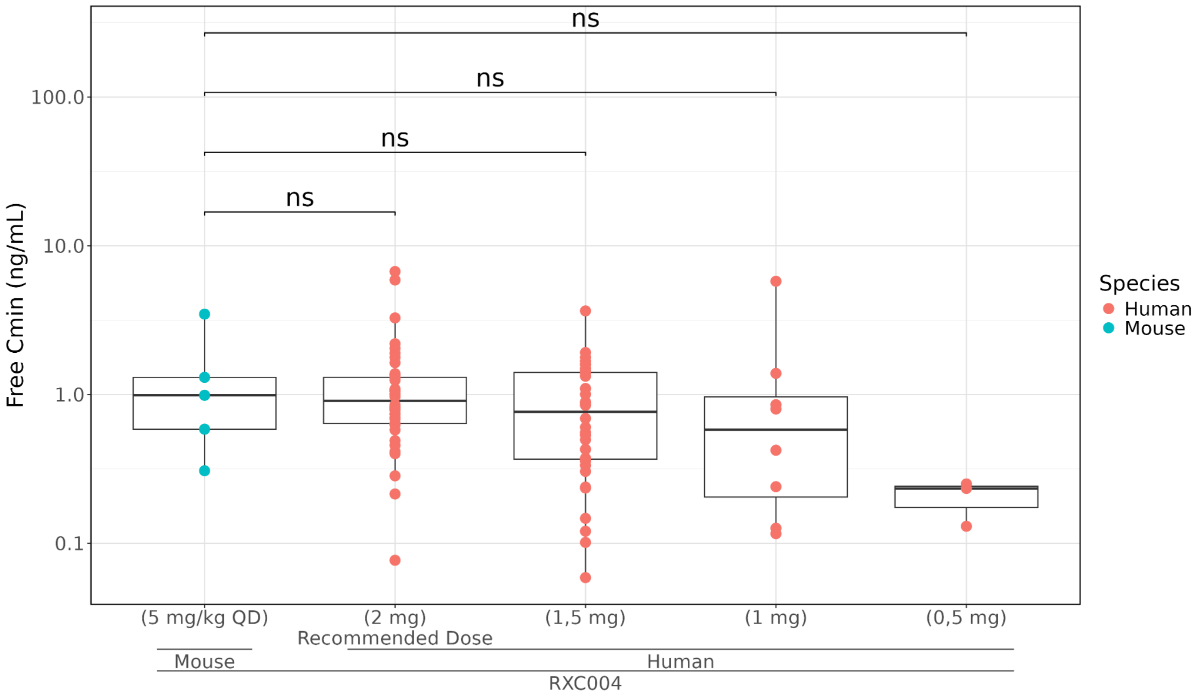


***Supplementary Figure 3: RXC004 drug exposure comparison in PDXs and humans.*** *Free Cmin comparison between Phase II patients at steady state (in red) at 2, 1.5, 1 and 0.5 mg and mice treated at 5 mg/kg QD (MED) after X days (in blue). One-tailed Wilcoxon p-values are reported after false discovery rate adjustment.*

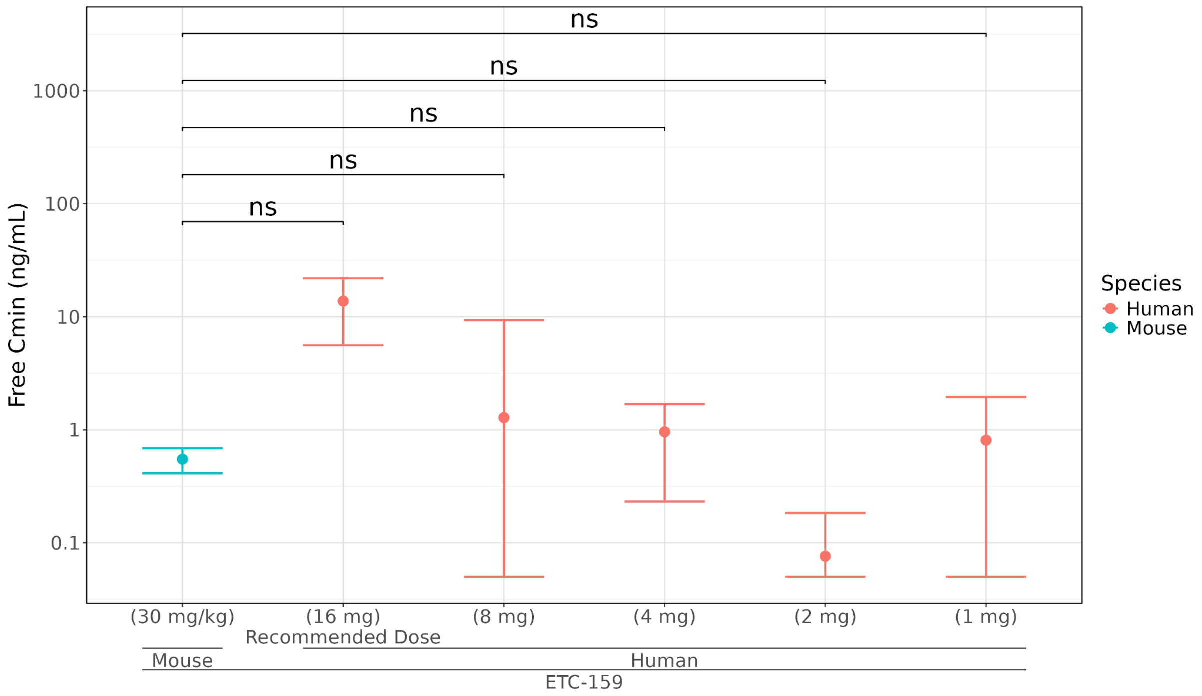


***Supplementary Figure 4: ETC159 drug exposure comparison in PDXs and humans.*** *Free Cmin comparison between Phase IA patients at steady state (C1D15) treated at 16, 8, 4, 2 and 1 mg QOD (in red) and mice treated after a single dose of ETC159 (in blue). One-tailed Welch p-values are reported after false discovery rate adjustment.*

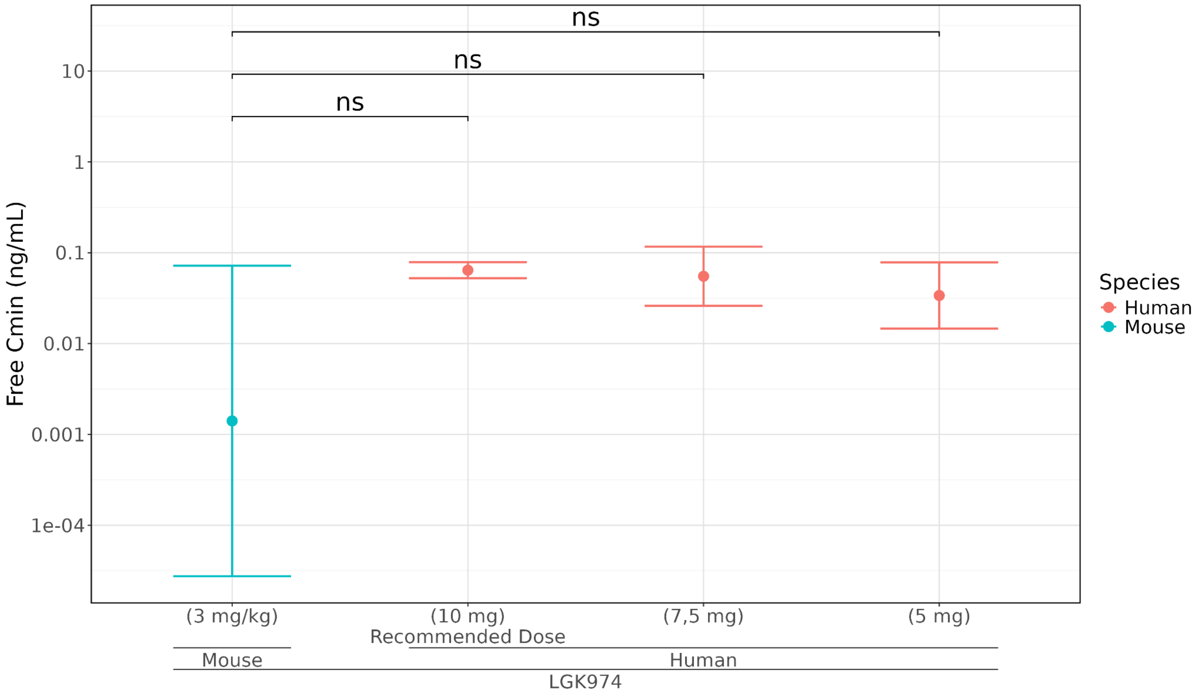


***Supplementary Figure 5: LGK974 drug exposure comparison in PDXs and humans.*** *Cmin comparison between Phase I patients at steady state (C1D15) treated at 10, 7.5, 5 and 2.5 mg QOD (in red) and nude mice treated after a single dose of LGK974 at 3 mg/kg (in blue). One-tailed Welch p-values are reported after false discovery rate adjustment.*

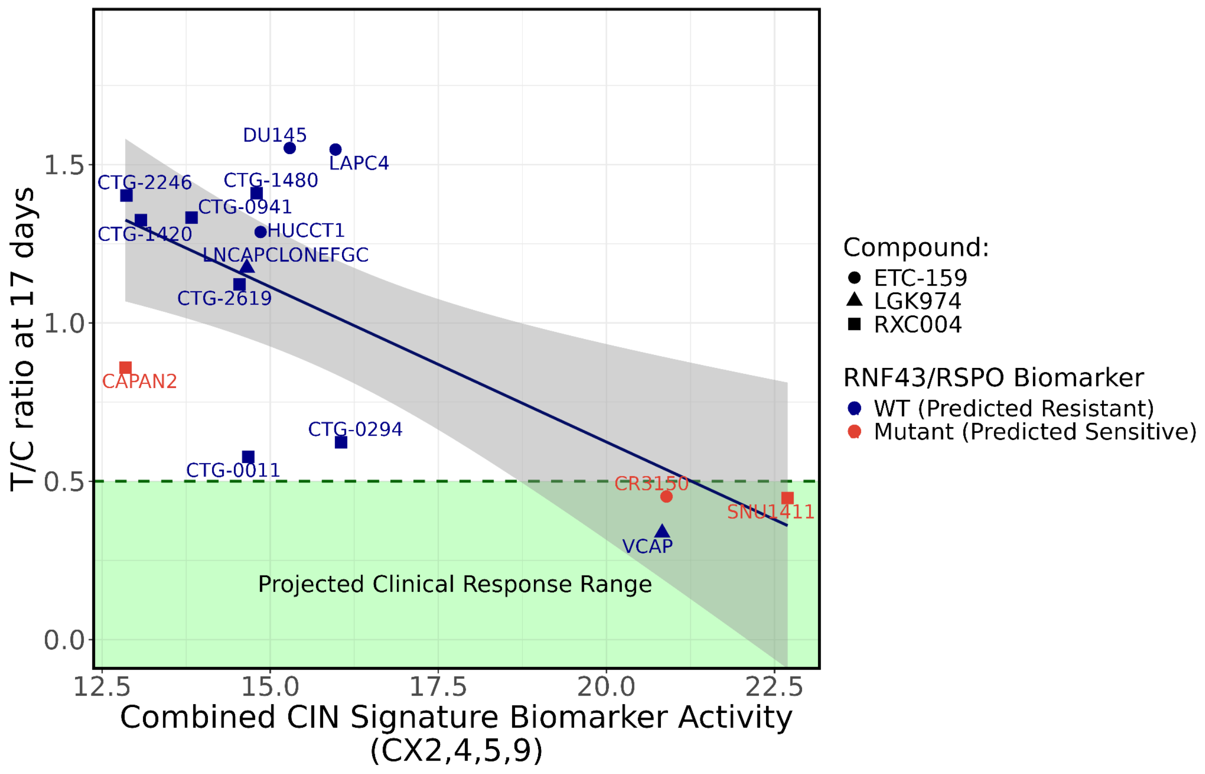


***Supplementary Figure 6: Comparison between RNF43 / RSPO Biomarker vs Combined CIN Signature Biomarker Activity.*** *RNF43 LoF/RSPO fusions do not result in consistent T/C ratios of less than 0.5, and therefore performs worse as a clinical biomarker for PORCN inhibitors. In comparison, high activity scores for the combined CIN signature biomarker see a T/C score of less than 0.5.*

| | Biliary Tract Cancer | Colorectal Adenocarcinoma | | Ovarian Cancer | Endometrial Cancer | Solid Tumours | Multiple Indications* | Cutaneous Melanoma |
|---|---|---|---|---|---|---|---|---|
| Trial | NCT04907851 | NCT04907539 | NCT02521844 | NCT02521844 | | NCT02521844 | NCT01351103 (dose escalation) | NCT01351103 (dose expansion) |
| Drug | RXC004 | RXC004 | ETC159 | ETC159 | | ETC159 | LGK974 | LGK974 |
| Observed Combination ORR | 0 (0 out of 14) | 0.286 (2 out of 7) | 0 (0 out of 10) | 0.222 (2 out of 9) | | 0.125 (1 out of 8) | 0.083 (1 out of 12) | 0.125 (5 out of 40) |
| Estimated Monotherapy ORR | 0 (0 out of 14) | 0.095 (1 out of 7) | 0 (0 out of 10) | 0.106 (1 out of 9) | | 0.033 (0 out of 8) | 0 (0 out of 12) | 0 (0 out of 40) |
| Projected Biomarker Prevalence | 0% | 2.9% | | 23.9% | 21.7% | 13.5% | 22.9% | 13.9% |
| Biomarker Predicted ORR (pORR) [95% CI] | 0 [0 - 0.232] | 0 [ 0 - 0.410] | 0 [0 - 0.308] | 0.222 [0.028 - 0.6] | | 0.125 [0.003 - 0.527] | 0.25 [0.055 - 0.572] | 0.125 [0.042 - 0.268] |
| Projected Responders [95% CI] | 0 [0 - 3] | 0 [0 - 2] | 0 [0 - 3] | 2 [0 - 5] | | 1 [0 - 4] | 3 [0 - 6] | 5 [1 - 10] |
| Probability > 30% ORR | 0 | 0 | 0 | 0.234 | | 0.014 | 0.564 | 0.017 |

***Supplementary Figure 7*. Summary efficacy data for PORCN inhibitors in combination with anti-PD-1 immunotherapy by cancer type and CIN signature prevalence.** *Across evaluated trials for RXC004, ETC159, and LGK974, biomarker-predicted ORRs closely mirrored observed combination outcomes, particularly in Cutaneous Melanoma and Ovarian Cancer cohorts. While indications with low biomarker prevalence (e.g., Biliary Tract and Colorectal cancers) consistently yielded 0% probability of achieving an ORR >30%, the Multiple Indications (LGK974) and Ovarian (ETC159) cohorts demonstrated the highest potential for enrichment, with predicted ORRs of 0.25 and 0.222 respectively. These data indicate that while observed combination efficacy remains modest, it aligns with predicted rates based on biomarker prevalence.*

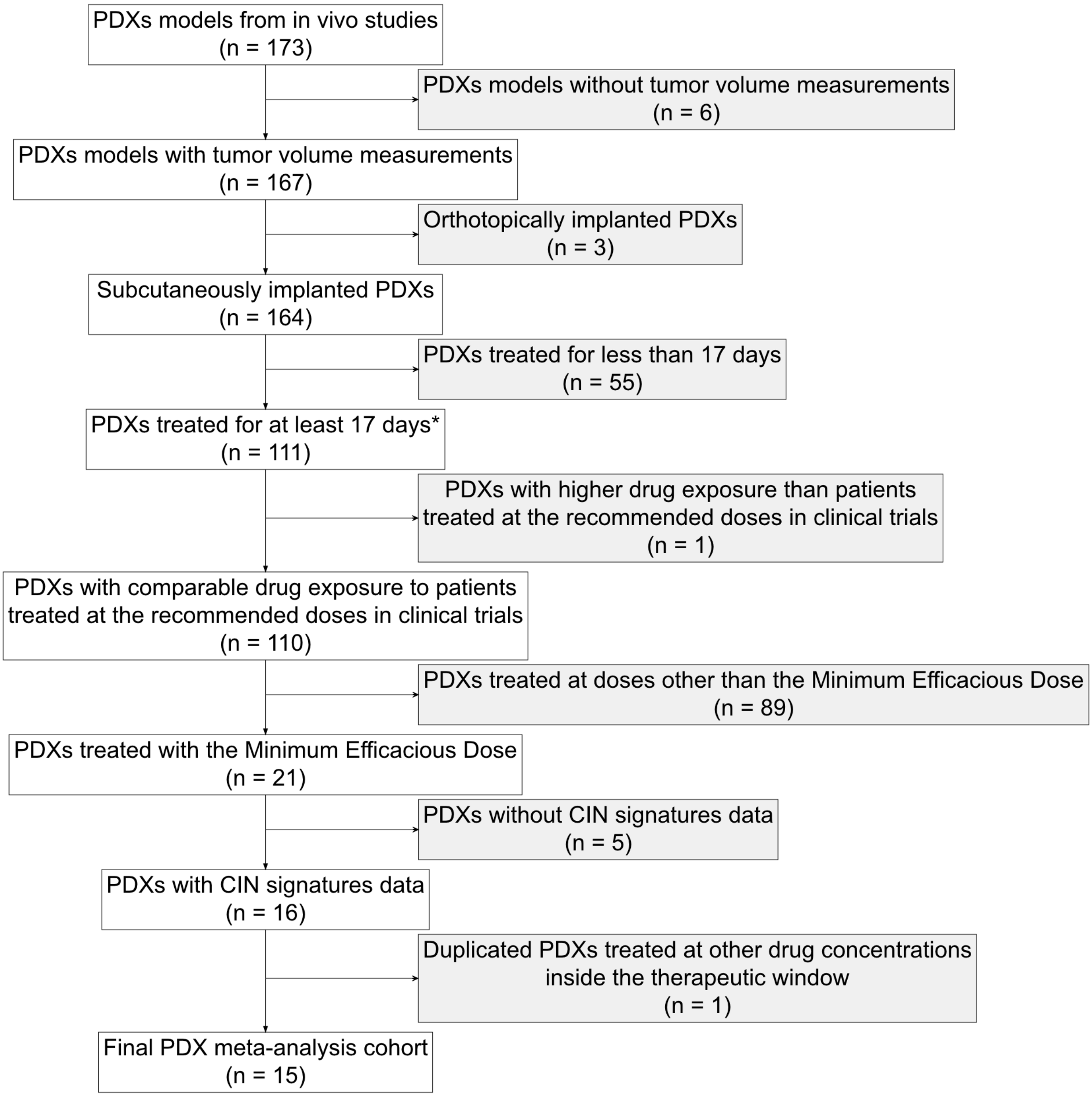


***Supplementary Figure 8 - Remark diagram of the PDX cohort.*** *REMARK, REporting recommendations for tumor MARKer prognostic studies. We identified 174 models that could potentially be added to our analysis. After exclusion of PDXs without tumor volume measurements (n=6) and orthotopically implanted (n=3) 165 models were eligible for analysis. T/C ratio measured at 17 days, was only available for 112 models (2 exceptions were made, HUCCT1 and DU145, see Methods section for details), from which only 22 were treated with the MED. CIN signatures were only available for 10 xenografts. After removing identical models treated at the same dose (n=1) we ended up with 9 xenografts in our PDX cohort.*

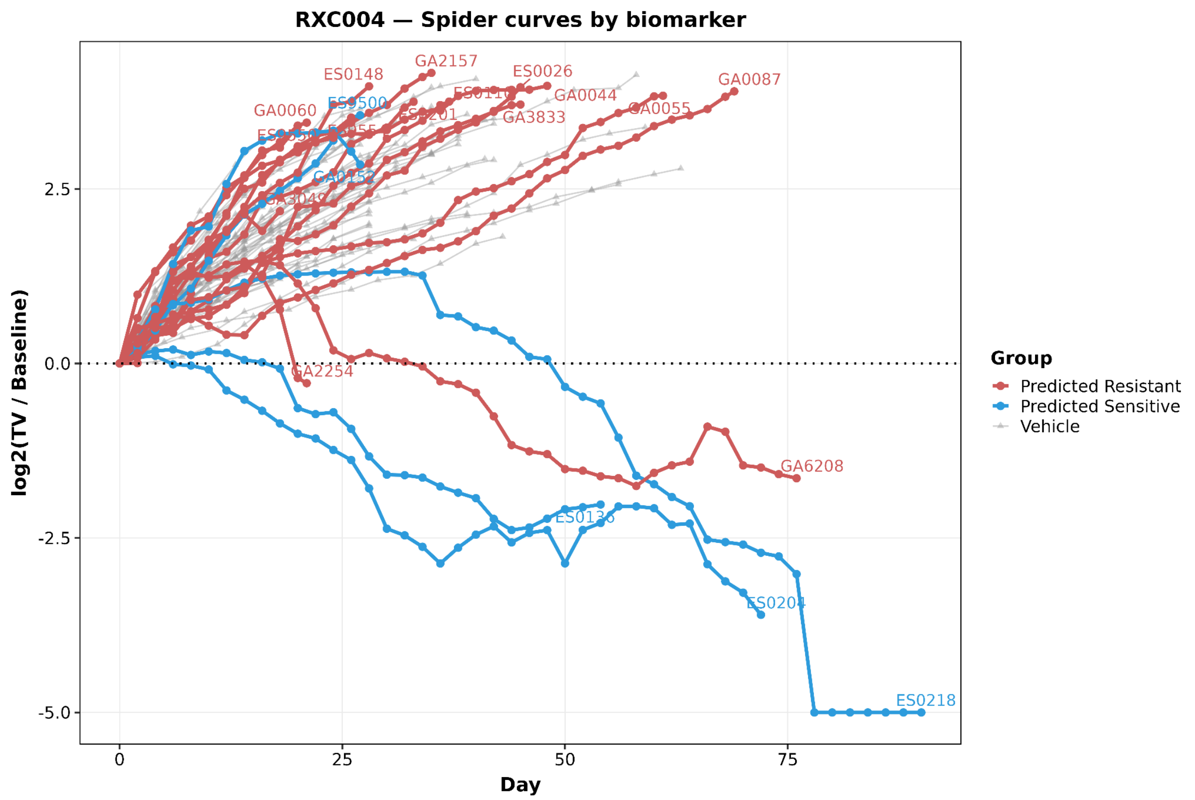


***Supplementary Figure 9 - Tumor volume curves of the PDXs enrolled in the prospective mouse clinical trial.*** *Log2 of the relative tumor volume at different time points for the PDXs mouse clinical trial. PDXs are colored based on their corresponding cohort: Combined CIN Signature Biomarker sensitive (blue) Combined CIN Signature Biomarker Sensitive resistant (red) and control group (gray)*

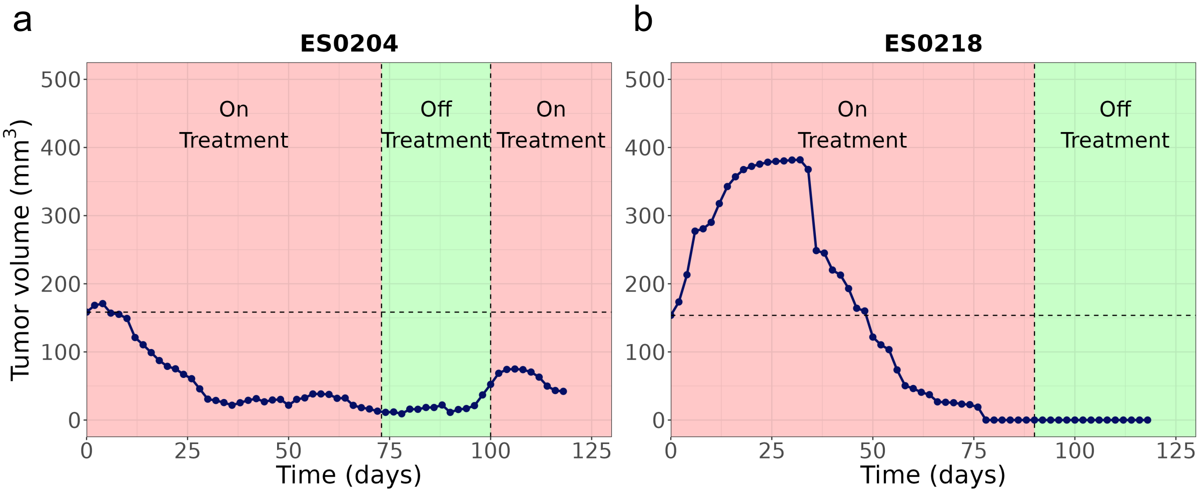


***Supplementary Figure 10 - Tumor volume curves of PDXs challenged on and off treatment.*** *ES0204 and ES0218 tumor volume curves in mm*$^3$ *showing the on treatment (red) and off treatment (green) periods for each model across time.*

#### Supplementary Table Captions

***Supplementary Table 1:*** *Results of combined CIN signature biomarker assessment.*

***Supplementary Table 2:*** *PDXs treated with PORCN inhibitors described in previous studies included in CIN biomarker association analysis.*

***Supplementary Table 3:*** *Free fraction in mice and humans for the different PORCNi.*

***Supplementary Table 4:*** *PDX models used for MED selection in mice.*

***Supplementary Table 5:*** *Mouse Cmin data used for the therapeutic window analysis. (*) Cmin measures were taken after single dose administration in mice.*

***Supplementary Table 6:*** *Human Cmin data used for the therapeutic window analysis. (*) Also includes 2 mg-2 weeks on/1 week off dosage regimen. (**) Also includes 1,5 mg+nivolumab and 1,5 mg+pembrolizumab. (***) Also includes 1mg+nivolumab. (****) Data represented as geometric mean instead of arithmetic mean.*

***Supplementary Table 7: PORCN inhibition response rates for every TCGA cancer type calculated based on the CIN signature threshold (t>19.33) obtained in the PDX analysis.*** *Some cancer types were created based upon the combining cohorts. COADREAD: colon adenocarcinoma + rectal adenocarcinoma, Gastric/Esophageal: Curated from a subset of Esophageal carcinoma + Stomach adenocarcinoma as per TCGA curation[79]. Solid tumours: All cancer types at the exception of leukemias.*

***Supplementary Table 8 - Anti-PD1 immunotherapy efficacy trials and associated 95% confidence interval around the ORR estimates.*** *Each row corresponds to a clincaltrials.org trial investigating the efficacy for an anti-PD1 immunotherapy (either Pembrolizumab or Nivolumab) used in the combination trials (Supplementary Figure 3). ORRs and DCRs extracted from these trials results and reported alongside calculated variance and lower bound of the 95% confidence interval of the ORR estimate (c.f. methods).*

***Supplementary Table 9 - RNF43 mutations and RSPO fusion in TCGA.*** *Predicted responders were defined by a CIN biomarker score > 19.33 threshold. RSPO3 fusions were identified via the GoF annotation in the TCGA (n=7,66, patients with microsatellite stable tumours only); RNF43 loss-of-function mutations were identified via the LoF annotation. No patient harboured both alterations. LoF, loss-of-function; OV, ovarian serous cystadenocarcinoma; STAD, stomach adenocarcinoma.*

***Supplementary Table 10:*** *PDX models assessed in the prospective mouse clinical trial.*
